# CLAMP directly interacts with MSL2 to facilitate *Drosophila* dosage compensation

**DOI:** 10.1101/415455

**Authors:** Evgeniya Tikhonova, Anna Fedotova, Artem Bonchuk, Vladic Mogila, Erica N. Larschan, Pavel Georgiev, Oksana Maksimenko

**Author notes:** Correspondence Oksana Maksimenko, Pavel Georgiev.

## Abstract

The binding of *Drosophila* male-specific lethal (MSL) dosage compensation complex exclusively to male X chromosome provides an excellent model system to understand mechanisms of selective recruitment of protein complexes to chromatin. Previous studies showed that the male-specific organizer of the complex, MSL2, and ubiquitous DNA-binding protein CLAMP are key players in the specificity of X chromosome binding. The CXC domain of MSL2 binds to genomic sites of MSL complex recruitment. Here we demonstrated that MSL2 directly interacts with the N-terminal zinc-finger domain of CLAMP. CLAMP-MSL2 and CXC-DNA interactions are cooperatively involved in recruitment of MSL complex to the X chromosome.

## Introduction

In *Drosophila*, a multisubunit dosage compensation complex (DCC) is responsible for an increase in the expression of genes on the X-chromosome in males (Kuroda et al, 2016; Lucchesi, 2018; Samata & Akhtar, 2018). One of the key questions is how the DCC specifically binds to the X-chromosome of males. This issue is of a fundamental importance for the understanding of the general mechanisms of specific binding of protein complexes to particular DNA sequences enmeshed in the chromatin *in vivo*.

The DCC consists of five proteins, MSL1, MSL2, MSL3, MOF, and MLE, and includes two non-coding RNAs, roX1 (3.7 kb) and roX2 (0.6 kb) that perform similar functions (Lucchesi, 2018; Samata & Akhtar, 2018). MSL1, MSL3, MOF, and MLE proteins are also present in females and are involved in the regulation of gene expression by a mechanism independent of the dosage compensation (Kuroda et al, 2016). MSL2 is expressed only in males suggesting a key role of this protein in binding specificity of DCC in males. The MSL1 and MSL2 proteins form a structural core of the DCC, while the C-terminal PEHE domain of MSL1 is responsible for the interaction with MSL3 and MOF (Hallacli et al, 2012; Kadlec et al, 2011; Li et al, 2005). The MLE protein, an ATP-dependent RNA/DNA helicase of the DEAD subfamily, interacts with two noncoding roX RNAs, roX1 and roX2, and induces their unwinding (Ilik et al, 2017; Maenner et al, 2013), as a result, making both roX RNAs capable of binding MSL2. Thus, roX RNAs link the MLE and MSL2 proteins into a single complex.

An incomplete MSL1-MSL2 complex binds a reproducible set of about 200 sites on the X chromosome, which were called the primary Chromatin Entry Sites (CES) (Alekseyenko et al, 2008) or High-Affinity Sites (HAS) (Straub et al, 2008). Within the HAS/CES, a GA-rich sequence motif, the MSL recognition element (MRE), is important for DCC targeting (Alekseyenko et al, 2008; Straub et al, 2008). Structural analysis showed that MSL2 specifically recognizes the MRE by its CXC domain (Fauth et al, 2010; Zheng et al, 2014). By using *in vitro* genome-wide DNA binding assays, Villa et al. (Villa et al, 2016) demonstrated that MSL2 specifically binds to a subclass of HAS named PionX (pioneering sites on the X). At the following step, roX, MLE and MSL3 are involved in spreading of DCC to nearby non-PionX HAS, and genes whose transcription should be elevated (Kuroda et al, 2016; Samata & Akhtar, 2018).

Recently, in a screen for transcription factors which determined the ability of the HAS to stimulate transcription in the S2 cells, CLAMP protein was identified which contains seven zinc finger domains of the C2H2 type (Larschan et al, 2012). It was shown that the C2H2 zinc fingers 4 through 7 of CLAMP specifically bind to the GA-repeats within the MREs (Kuzu et al, 2016; Soruco et al, 2013). A genome-wide ChIP-seq analysis has shown that CLAMP binds not only to HAS, but also to more than 5000 sites throughout the genome, most of which are distributed along the autosomes (Soruco et al, 2013). The latter finding complicates the understanding of the role of CLAMP in the specific recruitment of the DCC to the X-chromosome. Furthermore, it has been demonstrated that CLAMP opens chromatin at HAS which promotes MSL complex recruitment (Urban et al, 2017).

Despite performing a critical role in the MSL complex recruitment, immunostaining and ChIP sequencing (ChIP-seq) analysis reveals that CLAMP is not an X chromosome-specific protein and that only 3% of its binding sites overlap with those of the MSL complex (Soruco et al, 2013).

Here, we examined a possible direct role of CLAMP in recruiting the DCC. We tested the interactions of MSL proteins with CLAMP using the yeast two-hybrid assay and found that MSL2 directly interacts with CLAMP. A pull-down assay showed that the first zinc-finger C2H2 domain of CLAMP interacts with a highly conserved 620-655 aa region of MSL2. This interaction contributes to recruiting of the DCC to the X chromosome. Thus, MSL2 can recognize specific chromatin regions by additive interaction with DNA motif and CLAMP.

## Results and Discussion

### A conserved 36 aa domain of MSL2 directly interacts with the N-terminal zinc finger domain of CLAMP

Previous studies implicated CLAMP in recruitment of the DCC to the X chromosome (Soruco & Larschan, 2014). To identify direct interactions between CLAMP and components of the DCC, we used the yeast-two hybrid assay (Y2H). CLAMP (Fig.1A) contains a single zinc-finger C2H2 domain and a low complexity region at the N-terminus, and a cluster of six C2H2 domains at the C-terminus (Soruco et al, 2013). Y2H assay showed that CLAMP interacts with MSL2 and MLE proteins (data not shown).

Here, we examined the physical interactions between CLAMP and MSL2. The 773 aa MSL2 protein contains an N-terminal zinc RING finger (41-93 aa), a cysteine-rich CXC domain (520-567 aa), a C-terminal proline-rich (685-713 aa), and basic residue-rich (714-726 aa) regions (Fig. 1A, Appendix S1). We tested the interaction between CLAMP and MSL2 proteins using a pull-down assay *in vitro*. We tagged the 573-708 aa region of MSL2 and 1-196 aa of CLAMP with either GST or Thioredoxin-6xHis-tag. The GST protein and the 6xHis-tagged MSL2 RING domain (1-195 aa) were used as negative controls. Both GST- and 6xHis-tag pull-down assays confirmed interaction between 1-196 aa of CLAMP and 573-708 aa of MSL2 (Fig. 1B). In order to more precisely map the interacting domains, we developed a set of deletion derivatives of the GST-tagged MSL2 and Thioredoxin-6xHis-tagged CLAMP. The pull down experiments showed that a region between amino acids 86-153 of CLAMP containing the zinc-finger and 30 preceding residues interact with MSL2 (Fig. 1C), and amino acids 618-655 of MSL2 are critical for its interaction with CLAMP (Fig. 1D). The CLAMP interacting region in MSL2 is highly conserved within the *Drosophila* family, similar to the RING and CXC domains (Appendix S1). To further validate direct interactions of the proteins *in vivo*, we performed coimmunoprecipitation (co-IP) of the proteins that were transiently over-expressed in the *Drosophila* S2 cells (Fig. 1E). The over-expressed MSL2 was highly unstable in S2 cells, in contrast to the truncated derivatives of MSL2 lacking either the RING domain or the basic region. For this reason, we expressed a mutant variant of MSL2 (MSL2^ΔB^) lacking the basic region (714-726 aa) that was stable in the S2 cells (Appendix S2). We compared the co-IP of 3xHA-tagged CLAMP with the 3xFLAG-tagged MSL2^ΔB^ or MSL2^Δ13dΔB^ (Δ13d is deletion of the 642-654 aa) in S2 cells. 3xHA-CLAMP efficiently co-immunoprecipitated with MSL2^ΔB^ but not with MSL2^Δ13dΔB^ demonstrating that the 618-655 aa domain of MSL2 is required for interaction with CLAMP *in vivo*.

**Figure 1.**
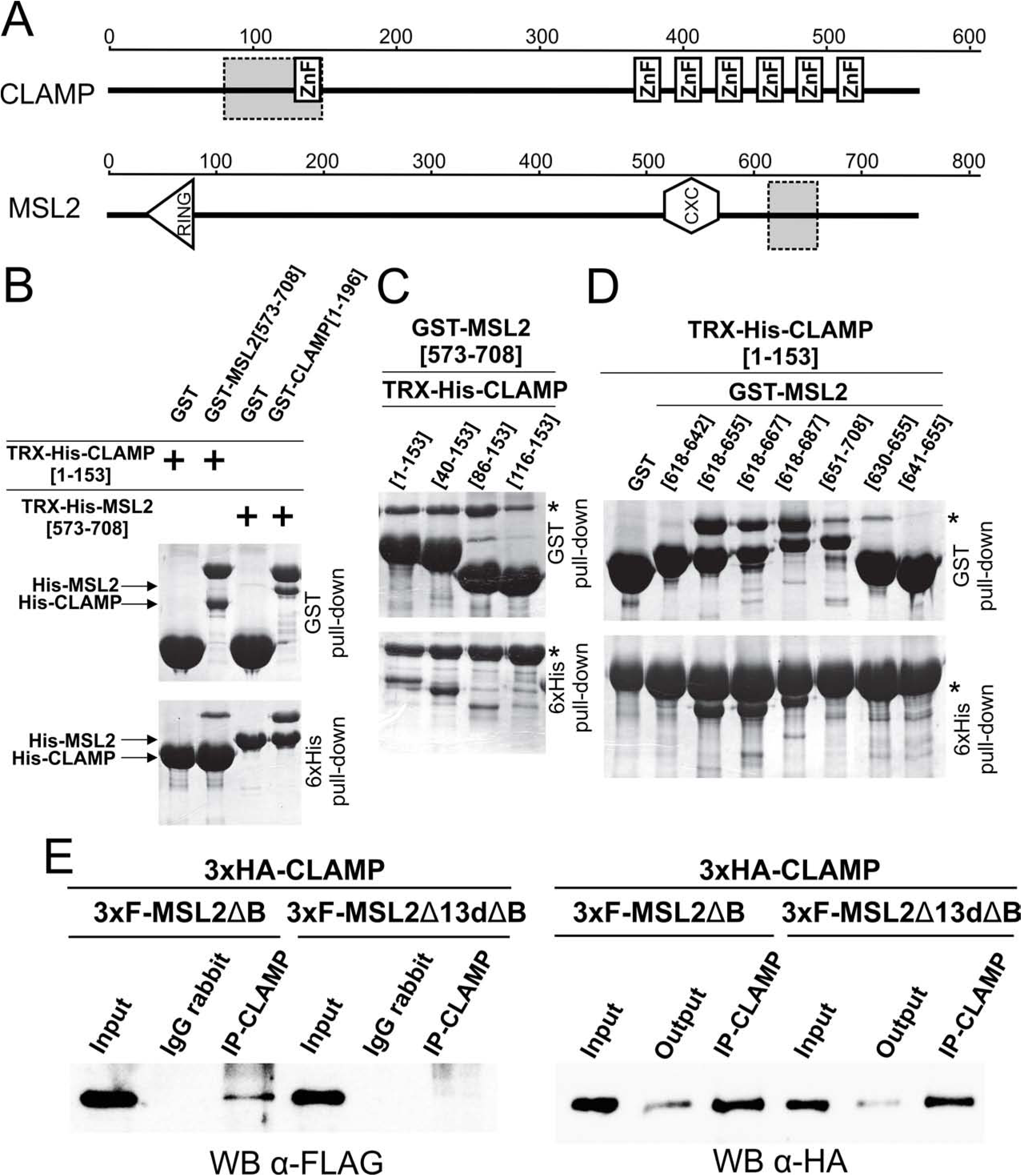
MSL2 directly interacts with CLAMP. **(A)** Schematic representation of full-length CLAMP and MSL2 proteins. The scale shows the number of amino acid residues. Boxes with dashed lines indicate interacting regions. **(B,C,D)** Confirmation of interacting domains of CLAMP and MSL2 proteins in vitro pull-down assay. **(B)** *In vitro* pull-down assay of either GST- or 6xHis-tagged MSL2[573-708] and CLAMP[1-196]. **(C)** Testing of the interaction *in vitro* between GST-MSL2[573-708] and Thioredoxin-6xHis-tagged deletion derivatives of CLAMP. The positions of amino acids are indicated by square brackets. Asterisk marks position of GST-MSL2[573-708]. **(D)** Testing of the interaction in vitro between Thioredoxin-6xHis-tagged CLAMP[1-153] and GST-tagged MSL2 deletion derivatives. Asterisk marks position of Thioredoxin-6xHis-CLAMP[1-153]. **(E)** Nuclear extracts from Drosophila S2 cells cotransfected with 3xHA-CLAMP and 3xMSL2 were immunoprecipitated with antibodies against CLAMP (using nonspecific rabbit IgG as a negative control), and the immunoprecipitates (IP) were analyzed by Western blotting for the presence of 3xFLAG-tagged MSL2 protein. The quality if immunoprecipitation was controlled by Western blotting for the presence of 3xHA-tagged CLAMP protein. Inputs show the starting samples of nuclear extract; outputs are supernatant after sedimentation of immunoprecipitated material.

### Inactivation of CXC or CLAMP-interacting region only partially affects MSL2 function in males

The MSL2 protein is critical for recruiting the DCC to the X chromosome in males (Lyman et al, 1997; Straub et al, 2013). It contains two well-characterized domains (Fig. 2A), the RING finger and CXC. In contrast, the functional roles of the MSL2 proline-rich (P) and basic (B) domains have not been characterized yet. There is some evidence that proline-rich and basic regions might be involved in interaction of MSL2 with the roX RNAs and binding of MSL2 to CES (Li et al, 2008; Villa et al, 2016).

**Figure 2.**
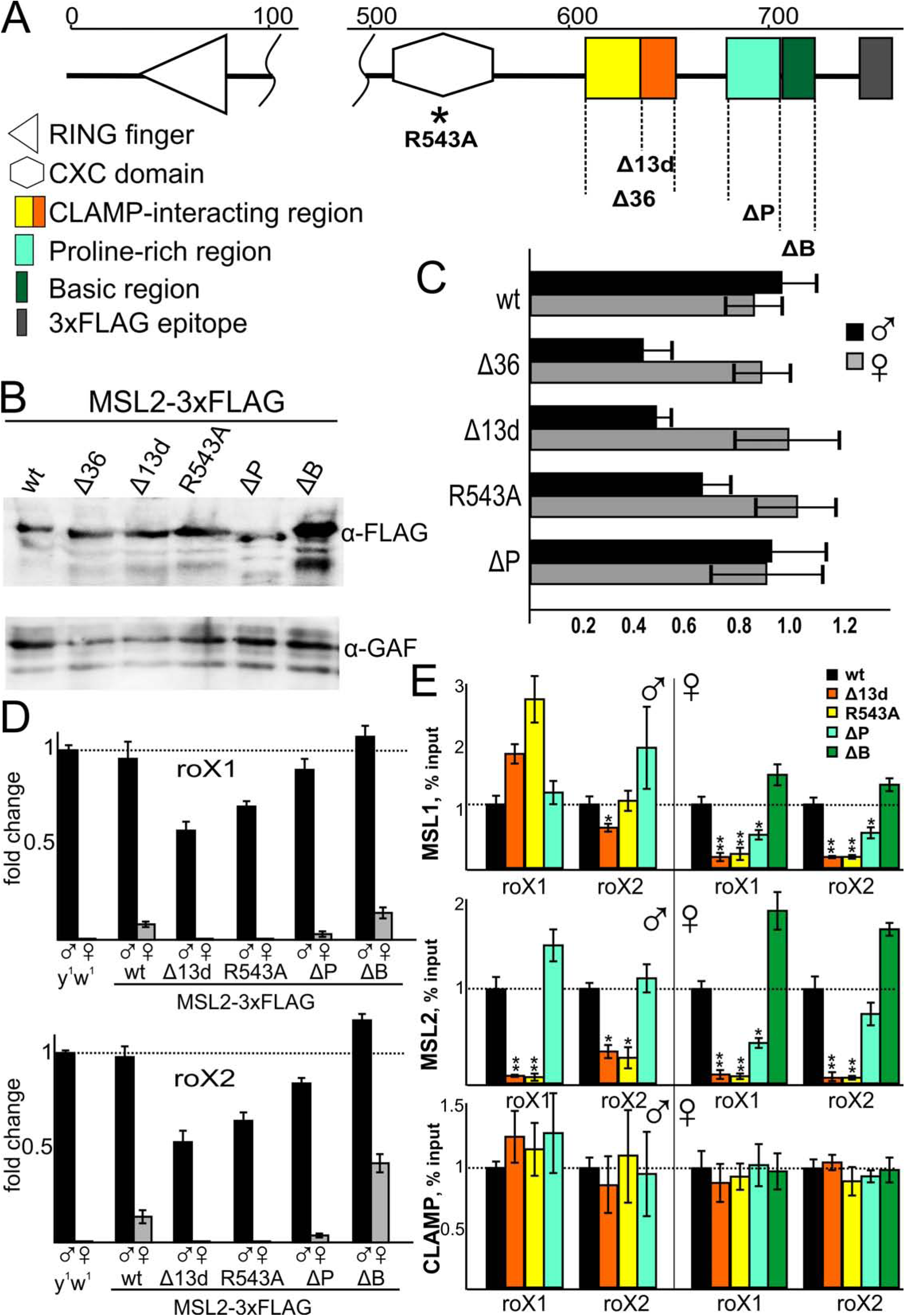
**(A)** Structure of MSL2 protein. The scale shows the number of amino acid residues. Dashed lines indicate deletions of full-sized MSL2 protein that used in this work. **(B)** Western blotting of protein extracts prepared from adult flies expressed different variants of 3xFLAG-tagged MSL2 protein. **(C)** Viability (as a relative percentage) of male (black) and female (gray) adult flies after expression of MSL2 derivatives at the msl2-null background (*msl2*^γ*227/γ227*^). Flies expressing corresponding MSL2 derivative in heterozygous *msl2*-null-mutant flies (*msl2*^γ*227*^/CyO) were used as internal controls with 100% viability. Results are expressed as mean with error bars show standard deviations of three independent crosses. **(D)** Expression levels of the roX1 and roX2 RNAs in male and female larvae in the *y*^*1*^*w*^*1*^ flies (wild-type background) and MSL2-expressing flies (wild-type (wt), deletion of 13 aa that interacts with CLAMP (Δ13d), substitution of arginine at the 543 position by alanine (R543A), deletion of proline-rich region (ΔP), deletion of basic region (ΔB)) at the *msl2*^γ*227*^ background. Individual transcript levels were determined by RT-qPCR with corresponding primers normalized relative to RpL32 for the amount of input cDNA. Histogram shows the changes of mRNAs for the tested *roX* genes comparing with expression levels in the wild type flies (corresponding to the *y*^*1*^*w*^*1*^ at the “1” on the scale). Error bars show standard deviations (n = 3). **(E)** Comparing binding of MSL1, MSL2 and CLAMP in the *roX1* and *roX2* CES regions in the MSL2-expressing flies at the *msl2*^γ*227*^ background. Histograms show ChIP enrichments at the CES regions on chromatin isolated from male and female larvae expressed different variants (wt, Δ13d, R543A, ΔP, ΔB) of MSL2 protein. The results are presented as a percentage of input genomic DNA normalized to corresponding positive autosomal genome regions for MSL1 (26E3), MSL2 (25A3), and CLAMP (39A1) and compared with binding levels in the flies with expressed wild type MSL2 protein (corresponding to the “1” on the scale). Error bars show standard deviations of quadruplicate PCR measurements for three independent experiments. Asterisks indicate significance levels: *p < 0.05, **p < 0.01.

To characterize domains of MSL2 that contribute to recruitment of the DCC to the X chromosome, we expressed in flies different cDNA variants of MSL2 tagged with 3xFlag at the C-terminus (Fig. 2A). To better define the CLAMP interacting region, we made a deletion not only of the whole 620-655 aa region (MSL2^Δ36^), but also of the distal 642-654 aa (MSL^Δ13d^). We also made MSL2 variants lacking either the proline-rich (Δ684-712, MSL2^ΔP^) or the basic (Δ714-727, MSL2^ΔB^) domain. To ablate activity of the CXC domain we used a previously characterized point mutation in the domain (Zheng et al, 2014) that substitutes the critical for DNA binding arginine at position 543 by alanine (R543A). A strong ubiquitin (Ubi) promoter was used to express the MSL2 variants. All transgenes were inserted at the same 86Fb region on the 3^rd^ chromosome using a φC31-based integration system (Bischof et al, 2007). Western blot analysis showed that all MSL2 variants were present in the transgenic flies at nearly equivalent levels (Fig.2B), with the exception of the MSL2^ΔB^ flies that expressed the FLAG-tagged MSL2 protein at a higher level than the others. Thus, MSL2^ΔB^ shows a higher stability than the wild type protein, both in flies and in the S2 cells.

Each transgene was introduced into the wild type and the null *msl2*^γ*227*^ background (Zhou et al, 1995). Ectopic expression of MSL2 in females results in assembling of functional DCC (Kelley et al, 1997) that leads to a strong elevation of gene expression, and as a consequence, to female lethality. As expected, females carrying homozygous MSL2^wt^ or MSL2^ΔB^ transgenes had a low viability (ca. 15% relative to males). Even heterozygous MSL2^wt^/+ or MSL2^ΔB^ /+ females had approximately 50% viability relative to males. In contrast, we observed normal viability in females carrying homozygous transgenes expressing mutant versions of MSL2 that affect either the CLAMP-interacting domain, the CXC domain, or the proline-rich domain. These results suggest that the functional activity of the latter transgenic MSL2 variants was at least partially compromised.

We also tested whether our mutant MSL2 transgenes can complement the null *msl2*^γ*227*^ mutation. Unexpectedly, all the mutant variants of MSL2 could complement *msl2*^γ*227*^ mutation and restored the viability in males homozygous for the null mutation. Viability was notably decreased approximately twofold only in males expressing the MSL2^Δ36^, MSL^Δ13d^, and MSL2^R543A^ transgenes in the *msl2*^γ*227*^ background (Fig.2C). These results suggest that all tested MSL2 variants are at least partially functional in their ability to recruit the DCC to the male X chromosome.

### CXC and CLAMP-interacting domains of MSL2 are critical for recruiting the DCC to the X-chromosome in females

In wild type female larvae, the roX RNAs are not expressed due to the absence of MSL complex, which activates their transcription (Meller et al, 1997). Expression of wild type MSL2 in females resulted in a significant activation of the roX2 RNA and to a lesser extent the roX1 RNA (Rattner & Meller, 2004). We confirmed that the expression of MSL2 in females induced transcription of the *roX* RNAs (Fig. 2D). However, expression of the roX RNAs in females was lower than in corresponding males. Expression of MSL2^ΔB^ in females resulted in an approximately twofold stronger expression of the roX RNAs, demonstrating a direct correlation between increased MSL2 concentration and activation of the *roX* genes. The HAS/CES located near the *roX* genes are required for stimulation of their transcription in males and repression in females (Bai et al, 2004; Urban et al, 2017). ChIP showed that MSL2 and MSL1 were bound to these sites in the MSL2^wt^, MSL2^ΔB^, and MSL2^ΔP^ transgenic female larvae, but the MSL2 mutants bearing the R543A or deletions of the CLAMP-interacting region lacked MSL1/MSL2 binding to these sites (Fig. 2E). These results demonstrate a direct correlation between the binding of MSL1/MSL2 and the activation of the *roX* genes in females. The only exception was the MSL2^ΔP^ transgene that still facilitated binding of MSL1/MSL2 to the HASs but did not stimulate *roX* expression. These results might suggest that the proline-rich region is involved in the activation of the *roX* genes.

In males, we observed only a relatively weak reduction (∼30-40%) of *roX* expression in the MSL2^Δ36^ and MSL2^R543A^ mutants. The wild type variant of MSL2 almost completely recapitulates normal level of *roX* expression. While in the MSL2^Δ36^ and MSL2^R543A^ mutants, the binding of MSL2 to the HASs, that regulate the *roX* genes, was notably reduced, MSL1 enrichment at the same sites was similar to that in the wild type males, or even higher (Fig. 2E). Therefore, it is possible that MSL1 can bind to the *roX* HASs not only as an MSL complex (DCC) subunit, but also as a part of an alternative MOF-containing complex(es), such as NSL (Chlamydas et al, 2016).

Polytene chromosomes in female salivary gland nuclei is a useful model system to test protein domains that are essential for recruiting the MSL complex to the X chromosome (Li et al, 2008; Morra et al, 2011; Zhou et al, 1995). In our experiments, the binding pattern was determined in at least five independent females from each transgenic line expressing MSL2 variants. For MSL2^wt^ and MSL2^ΔB^, we studied salivary glands in the animals heterozygous for the constructs. We used anti-FLAG mouse antibodies to identify targeted MSL2 variants, and anti-MSL1 rabbit antibodies to confirm recruitment of endogenous components of the MSL complex (Fig. 3, EV1). As a control, we also tested binding of the FLAG-tagged MSL2 variants to the polytene chromosomes in males expressing the endogenous MSL2 protein (Fig. EV1).

**Figure 3.**
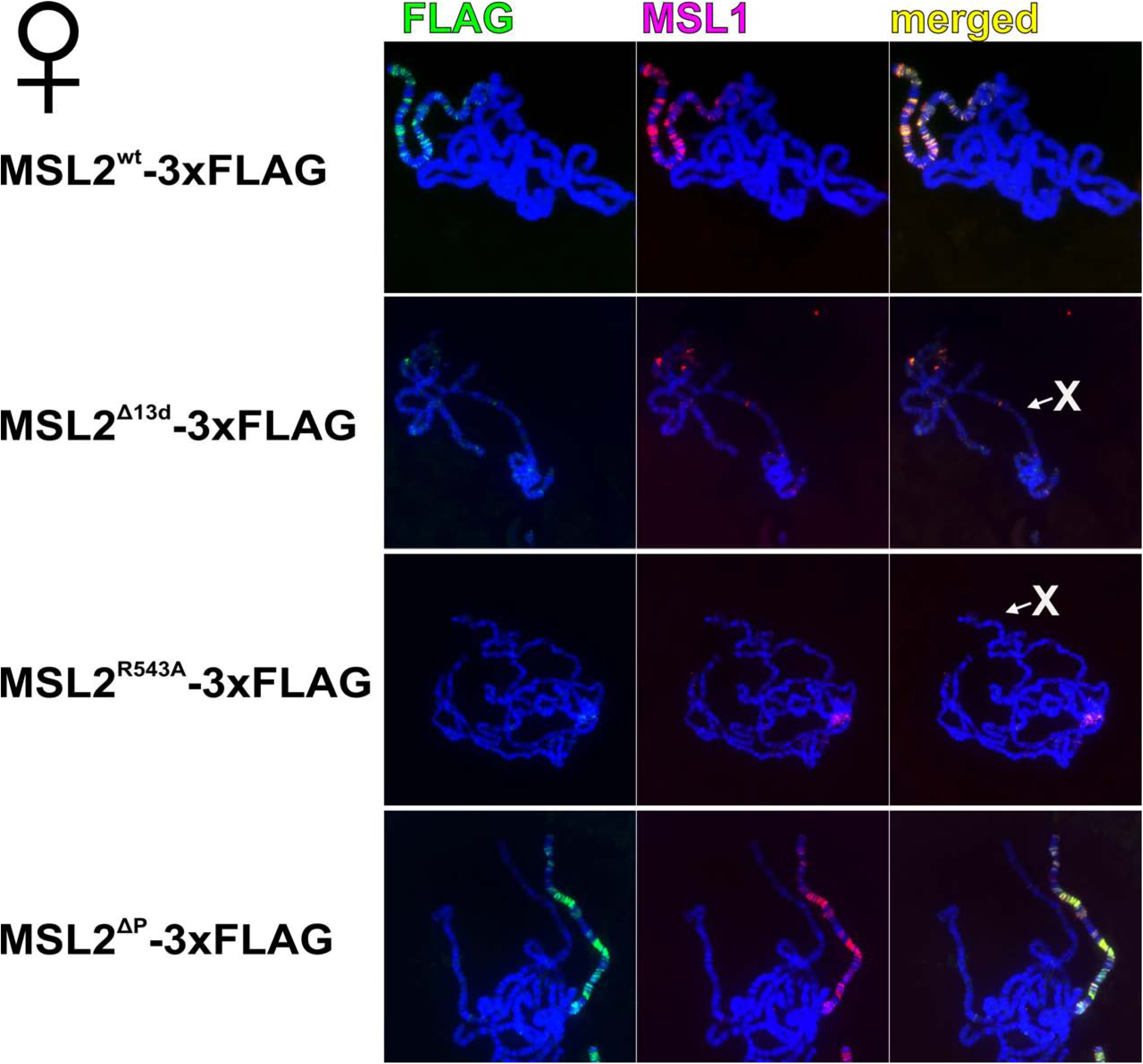
Distribution of MSL complex on the polytene chromosomes extracted from salivary glands of female 3^rd^ day larvae that express different FLAG-tagged variants of MSL2 protein: wild-type (wt), deletion of 13 aa from CLAMP-interacting region (Δ13d), substitution of arginine at the 543 position by alanine (R543A), deletion of proline-rich region (ΔP). Panels show immunostaining of 3xFLAG-MSL2 protein with mouse anti-FLAG antibody (green) and MSL1 protein with corresponding rabbit antibody (red) on polytene chromosomes. DNA is stained with DAPI (blue).

Transgenic expression of MSL2 in females led to localization to the X polytene chromosomes of both the MSL2 and MSL1 proteins, confirming that MSL2 expressed under the Ubi promoter is able to recruit MSL complex to the X chromosome in females (Fig. 3). The deletion of the basic domain did not affect the ability of the MSL2^ΔB^ mutant protein to recruit the DCC in females (Fig. EV1). The inactivation of the CXC domain in the MSL2^R543A^ transgenic larvae led to binding of the MSL2^R543A^ mutant to several sites on different chromosomes and the centromeric region (Fig. 3; Fig. EV1). MSL1 was co-localized at the same sites, suggesting that MSL2^R543A^ redirects recruitment of MSL complex to new sites. The deletions in the CLAMP interacting region (MSL2^Δ36^ and MSL2^Δ13d^) resulted in an almost complete absence of binding sites for the mutant variants of MSL2 and MSL1 suggesting an involvement of this domain in recruiting MSL complex. In addition, MSL2^ΔP^ and MSL1 were found at a limited number of sites (Fig. 3; Fig. EV1), which we identified as the high-affinity sites on the basis of the cytological map location for a selected subset (Lyman et al, 1997). This result is in agreement with the previous observation that roX RNAs are essential for recruitment and spreading of MSL complex from CES/HAS (Alekseyenko et al, 2008).

### The mutant MSL2 variants can support dosage compensation in males

All tested MSL2 mutants can, to a different extent, compensate for the null *msl2*^γ*227*^ mutation. Due to a strong disruption of MSL complex binding to the X-chromosome in females expressing the MSL2^Δ36^, MSL2^Δ13d^ or MSL2^R543A^ transgenes, it seems likely that strong expression of roX RNAs in males provides a compensatory mechanisms for recruiting MSL complex. We tested binding of the MSL complex to the polytene chromosome in the salivary gland nuclei of males carrying different variants of the MSL2 protein (Fig. 4, Appendix S3). In all cases, binding of MSL1 (rabbit antibodies) was correlated with the binding of the MSL2 variants (anti-FLAG mouse antibodies). MSL2^ΔP^ showed nearly the same binding pattern as MSL2^wt^ that correlates with the normal viability of MSL2^ΔP^ males. In contrast, in MSL2^R543A^ and MSL2^Δ13d^ male larvae, we observed reduced binding of MSL2-FLAG and MSL1 to the X chromosome (Fig. 4, Appendix S3). The same results were obtained with the anti-MSL2 rabbit antibodies (Fig. EV2). Additional MSL1/MSL2 bands were found on the polytene autosomes in the lines expressing MSL2^R543A^ and MSL2^Δ13d^, and to a lesser extent, in line expressing MSL2^ΔP^ (Fig. 4). This might be explained by the presence of unbound MSL complexes that bind to the lower affinity sites on autosomes.

**Figure 4.**
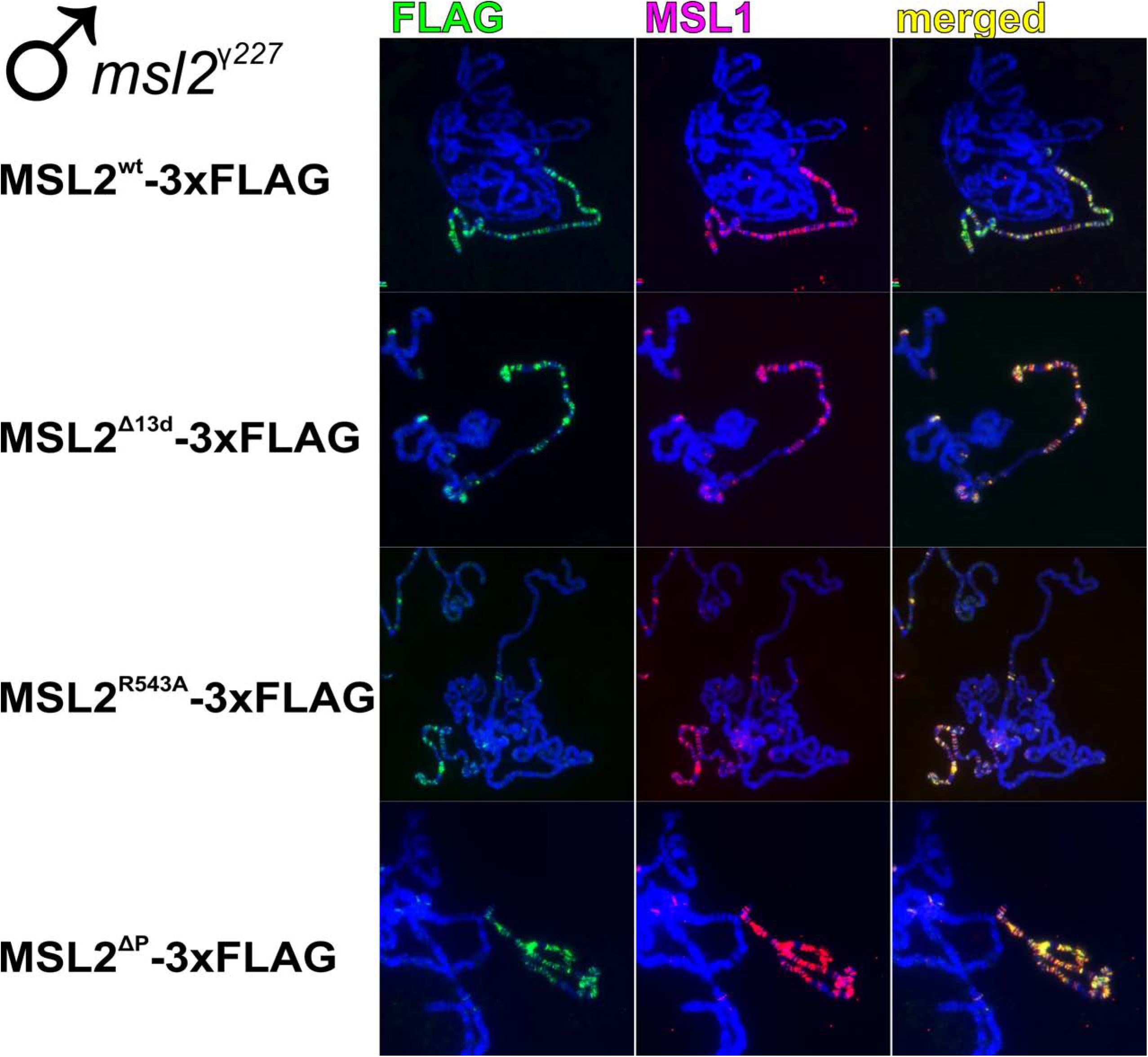
MSL1 and MSL2 localization on the polytene chromosomes extracted from salivary glands of male 3^rd^ day larvae at the *msl2*-null (*msl2*^γ*227*^) background that express different FLAG-tagged variants of MSL2 protein: wild-type (wt), deletion of 13 aa from CLAMP-interacting region (Δ13d), substitution of arginine at the 543 position by alanine (R543A), deletion of proline-rich region (ΔP). Panels show immunostaining of 3xFLAG-MSL2 protein with mouse anti-FLAG antibody (green) and MSL1 protein with corresponding rabbit antibody (red) on polytene chromosomes. DNA is stained with DAPI (blue).

To compare binding of MSL complex in different mutants, we selected previously characterized representative binding sites for MSL complex: PionX sites (Villa et al, 2016), CES sites (Alekseyenko et al, 2008), and several sites on the autosomes that have MSL2 signal (Straub et al, 2013). As a result, we found that MSL2-FLAG and MSL1 bound to all PionX and CES sites in males expressing wild type MSL2 (Fig. 5). MSL2^ΔP^ and MSL2^wt^ bound to all sites with the same or increased efficiency (Fig. EV3) supporting results obtained with the larval polytene chromosomes (Fig. 4). The MSL2^R543A^ and MSL2^Δ13d^ had the same binding patterns at all 16 tested sites, with no difference between PionX and CES (Fig. 5). At 12 sites, binding of MSL1 and the MSL2 mutant proteins was reduced nearly twofold. At 3 sites (2B14, 11B16, and 13D4) binding of both proteins did not change in mutant larvae, relative to control. Like in the case of the HAS regulated the *roX* genes, binding of the MSL2 mutant proteins, but not of MSL1, was reduced at the 9A3 site. At the same time, the binding of CLAMP was similar at all tested sites in the MSL2 mutants (Fig. EV4). We also observed that binding of the MSL1 (in MSL2^R543A^ and MSL2^Δ13d^ flies), but not the MSL2 mutant proteins was reduced at some autosomal sites (Fig.EV5). This result suggests an existence of alternative mechanisms of MSL2 recruitment to the sites outside of the DCC.

**Figure 5.**
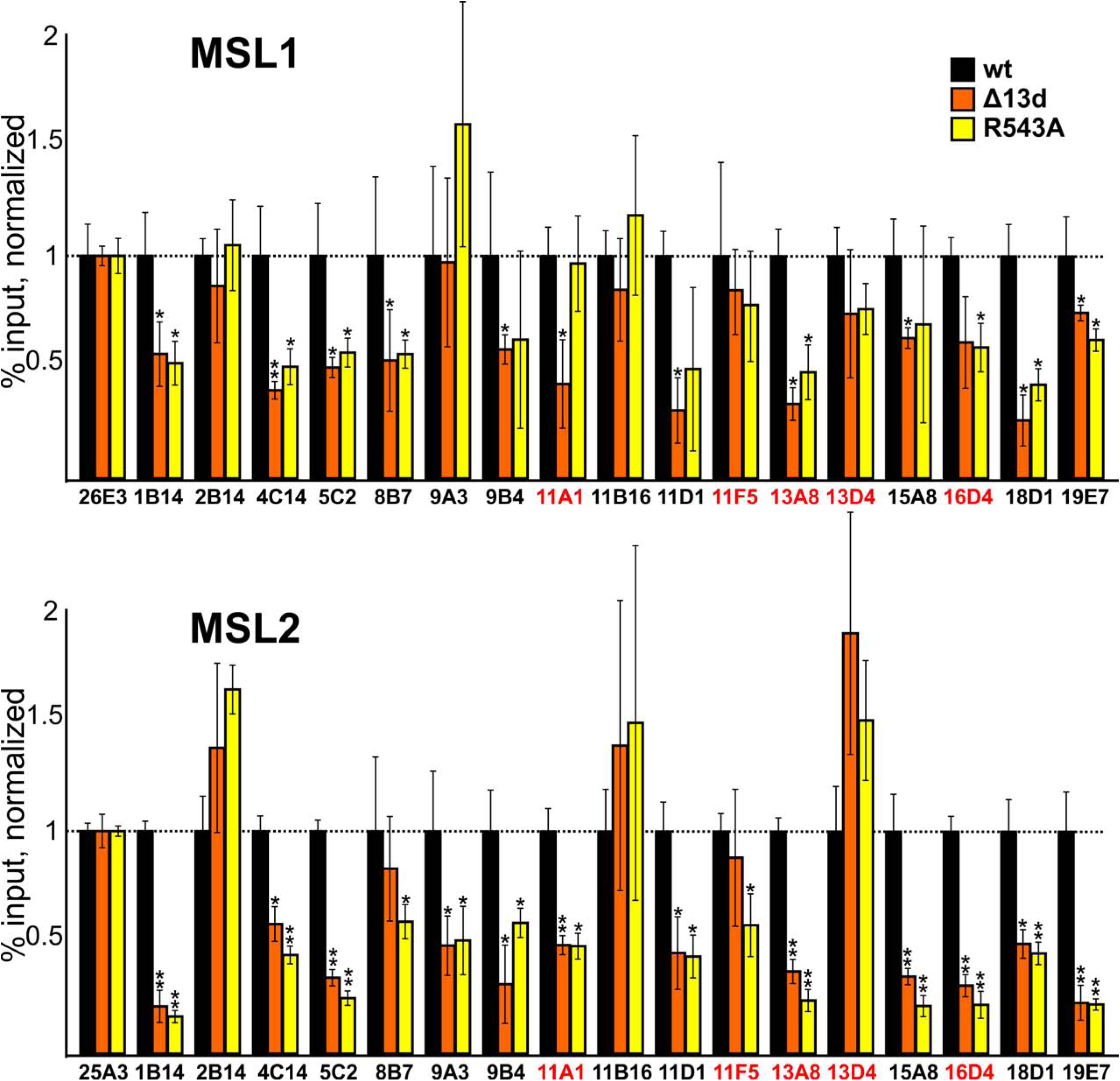
Comparing binding of MSL1 and MSL2 at the different CES and PionX (marked with red) regions in the MSL2-expressing flies at the *msl2*^γ*227*^ background. Histograms show ChIP enrichments at the CES regions on chromatin isolated from male flies expressed different variants (wt, Δ13d, R543A) of MSL2 protein. The results are presented as a percentage of input genomic DNA normalized to corresponding positive autosomal genome regions for MSL1(26E3) and MSL2 (25A3), and compared with binding levels in the flies with expressed wild type MSL2 protein (corresponding to the “1” on the scale). Error bars show standard deviations of quadruplicate PCR measurements for three independent experiments. Asterisks indicate significance levels: *p < 0.05, **p < 0.01.

Taken together, these results demonstrate that the previously characterized mutation in the CXC domain (MSL2^R543A^) and our new deletion of the CLAMP-interacting domain (MSL2^Δ13d^) alter recruitment of MSL complex to the X chromosome in the same way.

In conclusion, we found a direct interaction between a highly conserved region of MSL2 (618-655 aa) and the N-terminal C2H2 domain of CLAMP that binds to the CES/HAS regions. It seems likely that the MSL2 -- CLAMP interaction and the recognition of MRE by the CXC domain synergistically contribute to the specificity of the MSL complex recruitment to CES/HAS. The role of the CLAMP-interacting region is most obvious in females where MSL2/MSL1 binding to the *roX* promoters is critical for the expression of these RNAs. The CXC and CLAMP-interacting domains are essential for the initial recruitment of the MSL sub-complex (which likely also includes MSL1, MSL3, and MOF) to the regulatory regions of the *roX* genes. By connecting MLE and MSL2, roX RNAs induce the formation of the complete MSL complex. Previously, it was suggested that MLE is also involved in recruiting the MSL complex to the X chromosome (Straub et al, 2013). For this reason, the specific binding of MSL sub-complex (MSL1/MSL2) has become completely dependent on the CXC and CLAMP-interacting domains of MSL2. In males, the *roX* expression is less dependent on the binding of MSL complex (Bai et al, 2004; Meller et al, 1997). This might be a likely explanation of why the inactivation of the CXC or CLAMP-interacting domains in MSL2 only partially reduced recruitment of the MSL complex to the X chromosomes in males.

Furthermore, our results confirmed the previously described role of MSL2 in activating *roX* RNA transcription in females (Rattner & Meller, 2004). The proline-rich region of MSL2 might either directly contribute to activation of the *roX* promoters or block the CLAMP mediated silencing of the *roX* genes in females that was demonstrated recently (Urban et al, 2017). The basic region of MSL2 seems to be involved in regulation of MSL2 stability and contains potential sites for ubiquitination. As an E3 ligase, MSL2 promotes its autoubiquitination which is important for regulation of proper complex formation (Hallacli et al, 2012; Villa et al, 2012). Overall, our results are in agreement with the model that multiple specific protein-protein and protein-DNA interactions are responsible for recruitment of MSL complex to the X chromosome in males. Reduction of specificity of interactions with the X chromosome in lines expressing the MSL2 mutant variants leads to re-localization of the MSL complex to the autosomes and chromocenter that can interact with MSL complex with a lower affinity. Additional experiments are required to further support this model, e.g. by identification of new interactions between the components of the MSL complex and the proteins bound to CES/HAS.

## Materials and Methods

### Plasmid construction

For *in vitro* experiments protein fragments were either PCR-amplified using corresponding primers, or digested from MSL2 or CLAMP cDNA and subcloned into pGEX-4T1 (GE Healthcare) or into a vector derived from pACYC and pET28a(+) (Novagen) bearing p15A replication origin, Kanamycin resistance gene, and pET28a(+) MCS.

To express 3xFLAG-tagged MSL2 and 3xHA-tagged CLAMP in the S2 cells, protein-coding sequences were cloned in frame with 3×FLAG and 3xHA, excised, and subcloned into the pAc5.1 plasmid (Life Technologies). Plasmids for the yeast two-hybrid assay were prepared using the full-sized and truncated versions with separate domains of MSL2 and CLAMP as C-termini fused with pGBT9 and pGAD424 vectors from Clontech. PCR directed mutagenesis was used to make constructs with mutated MSL2 (corresponding primers are given in Appendix Table S1). Different full-sized variants of MSL2 were fused with 3xFLAG at the C-terminus and cloned into an expression vector. This vector contains *attB* site for φC31-mediated recombination, *Ubi67c* promoter with its 5’UTR, 3’UTR with SV40 polyadenylation signal, intronless *yellow* gene as a reporter for detection of transformants. Details of the cloning procedures, primers, and plasmids used for plasmid construction are available upon request.

### Pulldown assays

BL21 cells co-transformed with plasmids expressing GST-tagged and 6xHis-tagged derivatives of MSL2 and CLAMP were grown in LB media to an A600 of 1.0 at 37°C and then induced with 1 mM IPTG at 18°C overnight. ZnCl2 was added to final concentration 100 μM before induction. Cells were disrupted by sonication, centrifuged, applied to resin for 10 min at +4°C, followed by four washs with buffer C containing 500 mM NaCl. GST-pulldown was performed with Immobilized Glutathione Agarose (Pierce) in buffer C (20 mM HEPES-KOH, pH 7.7, 150 mM NaCl, 10 mM MgCl2, 0.1 mM ZnCl2, 0.1% NP40, 10% (w/w) Glycerol). The bound proteins were eluted with 50 mM reduced glutathione, 100 mM Tris-HCl, pH 8.0, 100 mM NaCl for 15 min. The 6xHis-pulldown was performed similarly, with Zn-IDA resin (Cube Biotech) in buffer A (50 mM HEPES-KOH, pH 7.6, with 500 mM NaCl, 5 mM MgCl2, 0.1 mM ZnCl2, 20 mM imidazole, 5% glycerol, 0.1% NP-40, and 5 mM β-mercaptoethanol) containing 1 mM PMSF and Calbiochem Complete Protease Inhibitor Cocktail VII (5 μL/mL), washed with buffer A containing 30 mM imidazole, and proteins were eluted with buffer B (50 mM HEPES-KOH, pH 7.6, with 500 mM NaCl, 250 mM imidazole, and 5 mM β-mercaptoethanol), 20 min at +4°C.

### Antibodies

Antibodies against MSL1[423-1030], MSL2[421-540], CLAMP[222-350] were raised in rabbits and purified from the sera by ammonium sulfate fractionation followed by affinity purification on CNBr-activated Sepharose (GE Healthcare, United States) or Aminolink Resin (ThermoFisher Scientific) according to standard protocols. Mouse monoclonal anti-FLAG M2 and anti-HA antibodies were from Sigma (Unites States).

### Cell culture, transfection

*Drosophila* S2 cells were grown in SFX medium (HyClone) at 25°C. Transfection of plasmids was performed with the Cellfectin II reagent (Invitrogen) according to the manufacturer’s instructions. Typically, cells were transfected in six-well plates and grown for 24 to 48 hours before harvesting.

### Co-immunoprecipitation assay

S2 cells grown in SFX medium were co-transfected by 3xFLAG-MSL2 (with deletions) and 3xHA-CLAMP plasmids with Cellfectin II (Life Technologies), as recommended by the manufacturer. After transfection, the cells were incubated for 48 h and then collected by centrifugation at 700 *g* for 5 min, washed once with 1×PBS and resuspended in 20 packed cell volumes of hypotonic lysis buffer (20 mM Tris-HCl, pH 7.4, with 10 mM KCl, 10 mM MgCl2, 2 mM EDTA, 10% glycerol, 1% Triton X-100, 1 mM DTT, and Calbiochem Complete Protease Inhibitor Cocktail V. After incubation on ice for 10 min, the cells were sonicated (2 x 15 s on ice at 20% output, 40% duty cycle), NaCl was added to a final concentration of 420 mM, and incubation on ice continued for 60 min, with periodic mixing. Sonication was repeated as above to reduce viscosity, cell debris was pelleted by centrifugation at 10,000 *g* for 30 min at 4°C, and the supernatant was collected for immunoprecipitation with anti-CLAMP antibodies, incubated with Protein A Agarose (Pierce), and equilibrated with incubation buffer-150 (20 mM Tris-HCl, pH 7.4, 150 mM NaCl, 10 mM MgCl2, 1 mM EDTA, 1 mM EGTA, 10% glycerol, and 0.1% NP-40). Rabbit IgG conjugated to Protein A Agarose beads (by incubating in the same buffer on a rotary shaker at 4°C for 1 h) were used as negative control. The protein extract (50 μg protein) was adjusted to a volume of 500 μL with buffer-150, mixed with antibody-conjugated beads (30 μL), and incubated on a rotary shaker overnight at 4°C. The beads were then washed with one portion of incubation buffer-500 (with 500 mM NaCl), one portion of incubation buffer-300 (with 300 mM NaCl), and one portion of incubation buffer-150, resuspended in SDS-PAGE loading buffer, boiled, and analyzed by Western blotting. Proteins were detected using the ECL Plus Western Blotting substrate (Pierce) with anti-FLAG and anti-HA antibodies.

### Fly crosses and transgenic lines

*Drosophila* strains were grown at 25°C under standard culture conditions. The transgenic constructs were injected into preblastoderm embryos using the φC31-mediated site-specific integration system at locus 86Fb (Bischof et al, 2007). The emerging adults were crossed with the *y ac w*^*1118*^ flies, and the progeny carrying the transgene in the 86Fb region were identified by *y*^*+*^ pigmented cuticle. Details of the crosses and primers used for genetic analysis are available upon request.

### Fly extract preparation

20 adult flies were homogenized with a pestle in 200 μL of 1xPBS containing 1% β-mercaptoethanol, 10 mM PMSF, and 1:100 Calbiochem Complete Protease Inhibitor Cocktail VII. Suspension was sonicated 3 times for 5 s at 5 W. Then, 200 μL of 4xSDS-PAGE sample buffer was added and mixture was incubated for 10 min at 100°C and centrifuged at 16,000 *g* for 10 min.

### *RNA isolation and* quantitative analysis

Total RNA was isolated from 2- to 3-day-old adult males and females using the TRI reagent (Molecular Research Center, United States) according to the manufacturer’s instructions. RNA was treated with two units of Turbo DNase I (Ambion) for 30 min at 37°C to eliminate genomic DNA. The synthesis of cDNA was performed using 2 μg of RNA, 50 U of ArrayScript reverse transcriptase (Ambion), and 1 μM of oligo(dT) as a primer. The amounts of specific cDNA fragments were quantified by real-time PCR with Taqman probes. At least three independent measurements were made for each RNA sample. Relative levels of mRNA expression were calculated in the linear amplification range by calibration to a standard genomic DNA curve to account for differences in primer efficiencies. Individual expression values were normalized with reference to *RpL32* mRNA.

The sequences of primers and oligonucleotides used in this work are given in Appendix Table S1.

### Immunostaining of polytene chromosomes

*Drosophila* 3^rd^ instar larvae were cultured at 18°C under standard conditions. Polytene chromosome staining was performed as described (Murawska & Brehm, 2012). The following primary antibodies were used: rabbit anti-MSl1at 1:100 dilution, rabbit anti-Msl2 at 1:100 dilution, and monoclonal mouse anti-FLAG at 1:100 dilution. The secondary antibodies were Alexa Fluor 488 goat anti-mouse 1:2,000 and Alexa Fluor 555 goat anti-rabbit 1:2,000 (Invitrogen). The polytene chromosomes were co-stained with DAPI (AppliChem). Images were acquired on the Nikon Elclipse T*i* fluorescent microscope using Nikon DS-Qi2 digital camera, processed with ImageJ 1.50c4 and Fiji bundle 2.0.0-rc-46. 3-4 independent staining and 4-5 samples of polytene chromosomes were performed with each MSL2-expressing transgenic line.

### Chromatin immunoprecipitation

Chromatin preparation was performed as described (Maksimenko et al, 2015; Zolotarev et al, 2017) with some modifications. Samples of 500 mg each of larva or adult flies were ground in a mortar in liquid nitrogen and resuspended in 10 mL of buffer A (15 mM HEPES-KOH, pH 7.6, 60 mM KCl, 15 mM NaCl, 13 mM EDTA, 0.1 mM EGTA, 0.15 mM spermine, 0.5 mM spermidine, 0.5% NP-40, 0.5 mM DTT) supplemented with 0.5 mM PMSF and Calbiochem Complete Protease Inhibitor Cocktail V. The suspension was then homogenized in a Dounce homogenizer with tight pestle and filtered through 50 μm Nylon Cell Strainer (BD Biosciences, United States). The homogenate was transferred to 3 mL of buffer A with 10% sucrose (AS), and the nuclei were pelleted by centrifugation at 4,000 g, 4°C, for 5 min. The pellet was resuspended in 5 mL of buffer A, homogenized again in a Dounce homogenizer, and transferred to 1.5 mL of buffer AS to collect the nuclei by centrifugation. The nuclear pellet was resuspended in wash buffer (15 mM HEPES-KOH, pH 7.6, 60 mM KCl, 15 mM NaCl, 1 mM EDTA, 0.1 mM EGTA, 0.1% NP-40, Calbiochem Complete Protease Inhibitor Cocktail V) and cross-linked with 1% formaldehyde for 15 min at room temperature. Cross-linking was stopped by adding glycine to a final concentration of 125 mM. The nuclei were washed with three 10 mL portions of wash buffer and resuspended in 1.5 mL of nuclear lysis buffer (15 mM HEPES, pH 7.6, 140 mM NaCl, 1 mM EDTA, 0.1 mM EGTA, 1% Triton X-100, 0.5 mM DTT, 0.1% sodium deoxycholate, 0.1% SDS, Calbiochem Complete Protease Inhibitor Cocktail V). The suspension was sonicated (20×30 s with 60 s intervals, on ice at 50% output), and 50 μL aliquots were used to test the extent of sonication and to measure DNA concentration.

Debris was removed by centrifugation at 14,000 *g*, 4°C, for 10 min, and chromatin was pre-cleared with Protein A agarose (Pierce), blocked with BSA and salmon sperm DNA, with 50 μL aliquots of such pre-cleared chromatin being stored as input material. Samples containing 10–20 μg of DNA equivalent in 1 mL of nuclear lysis buffer were incubated overnight, at 4°C, with rabbit antibodies against MSL1 (1:500), MSL2 (1:200), and CLAMP (1:200), or with nonspecific IgG purified from rabbit preimmune sera (control). Chromatin–antibody complexes were collected using blocked Protein A agarose at 4°C over 5 h. After several rounds of washing with lysis buffer (as such and with 500 mM NaCl), and TE buffer (10 mM Tris-HCl, pH 8; 1 mM EDTA), the DNA was eluted with elution buffer (50 mM Tris-HCl, pH 8.0; 1 mM EDTA, 1% SDS), the cross-links were reversed, and the precipitated DNA was extracted with phenol– chloroform. The enrichment of specific DNA fragments was analyzed by real-time PCR using a QuantStudio 12K Flex Cycler (Applied Biosystems). The primers used for PCR in ChIP experiments for genome fragments are shown in Appendix Table S1.

At least three independent biological replicates were made for each chromatin sample. The results of chromatin immunoprecipitation are presented as a percentage of input genomic DNA normalized to a positive control genomic site. Error bars show standard deviations of quadruplicate PCR measurements for three independent experiments. The *tubulin-γ37C* coding region (devoid of binding sites for the test proteins) were used as negative controls; Autosomal MSL1-binding region 26E3, MSL2-binding region 25A3, and CLAMP-binding region 39A1 were used as positive genomic controls.

## Acknowledgments

We thank Farhod Hasanov and Aleksander Parshikov for fly injections, and Olga Kyrchanova for help with fly genetics. This study was supported by the Russian Science Foundation, project no. 17-74-20155 (to O.M.). This study was performed using the equipment of the IGB RAS Core Facilities Centre, supported by the Ministry of Science and Education of the Russian Federation.

## Author contributions

PG, ENL and OM conceived the study; EA performed work with transgenic lines and coimmunoprecipitation experiments; AF and VM performed immunostaining of polytene chromosomes; AB performed pull-down assay; OM performed and analyzed RoX levels and ChIP experiments; All authors analyzed data; PG and OM provided supervision; ENL, PG and OM wrote the manuscript; OM provided funding.

## Expanded View Figure Legends

**Figure EV1.**
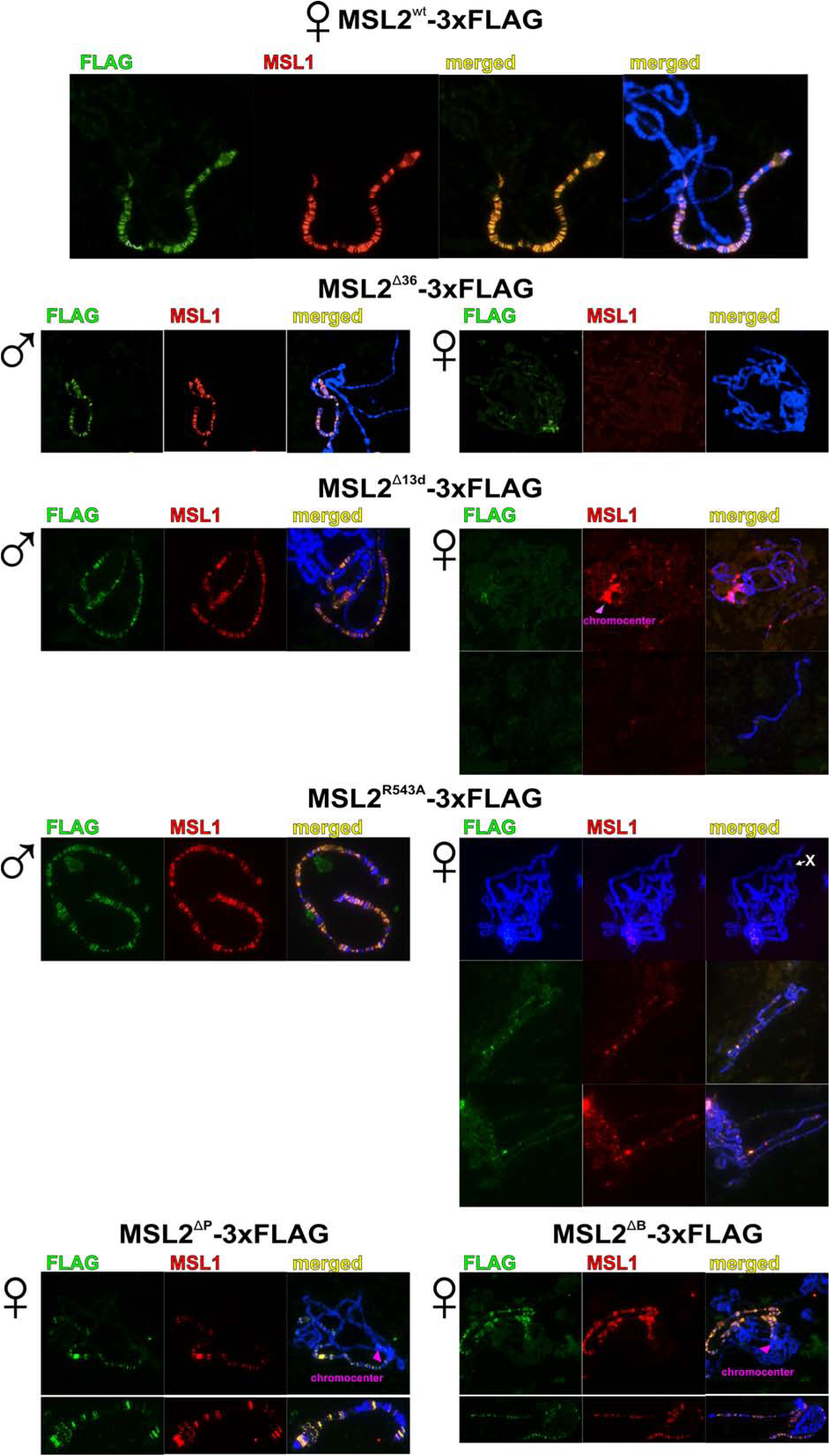
Distribution of MSL complex on the polytene chromosomes extracted from salivary glands of male and female 3^rd^ day larvae that express different FLAG-tagged variants of MSL2 protein: wild-type (wt), deletion of 13 aa from CLAMP-interacting region (Δ13d), substitution of arginine at the 543 position by alanine (R543A), deletion of proline-rich region (ΔP), deletion of basic region (ΔB). Panels show immunostaining of 3xFLAG-MSL2 protein with mouse anti-FLAG antibody (green) and MSL1 protein with corresponding rabbit antibody (red) on polytene chromosomes. DNA is stained with DAPI (blue).

**Figure EV2.**
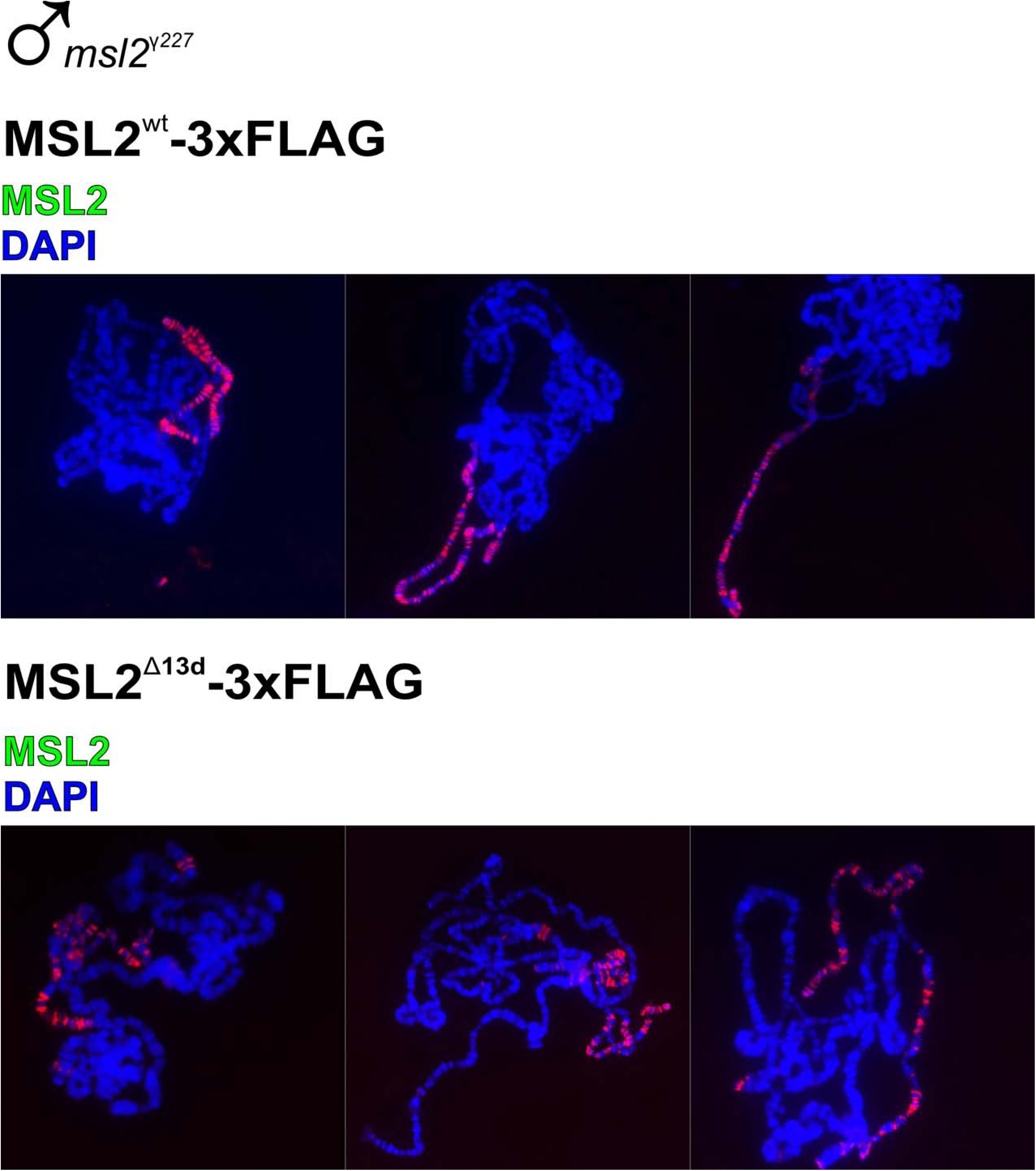
Immunostaining of polytene chromosomes extracted from salivary glands of male 3^rd^ day larvae at the *msl2*-null (*msl2*^γ *227*^) background that express FLAG-tagged variants of MSL2 protein: wild-type (wt) and with deletion of 13 aa from CLAMP-interacting region (Δ13d). Panels show immunostaining of 3xFLAG-MSL2 protein with rabbit anti-MSL2 antibodies (green) on polytene chromosomes. DNA is stained with DAPI (blue).

**Figure EV3.**
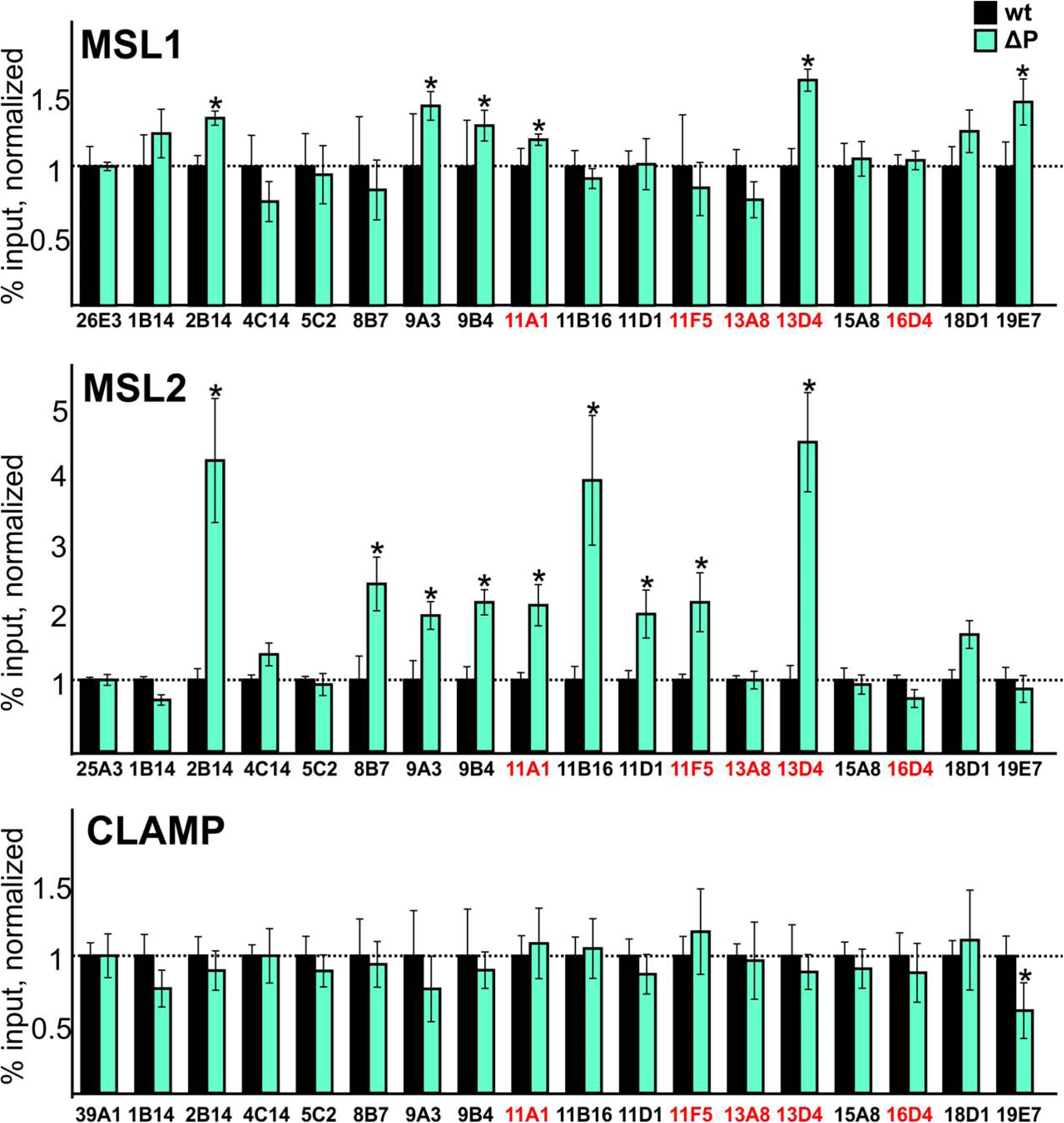
Comparing binding of MSL1, MSL2 and CLAMP at the different CES and PionX (marked with red) regions in the MSL2-expressing flies at the *msl2*^γ *227*^ background. Histograms show comparison of ChIP enrichments at the CES regions on chromatin isolated from male flies expressed wt and ΔP variants of MSL2 protein. The results are presented as a percentage of input genomic DNA normalized to corresponding positive autosomal genome regions for MSL1(26E3), MSL2 (25A3) and CLAMP (39A1), and compared with binding levels in the flies with expressed wild type MSL2 protein (corresponding to the “1” on the scale). Error bars show standard deviations of quadruplicate PCR measurements for three independent experiments. Asterisks indicate significance levels: *p < 0.05.

**Figure EV4.**
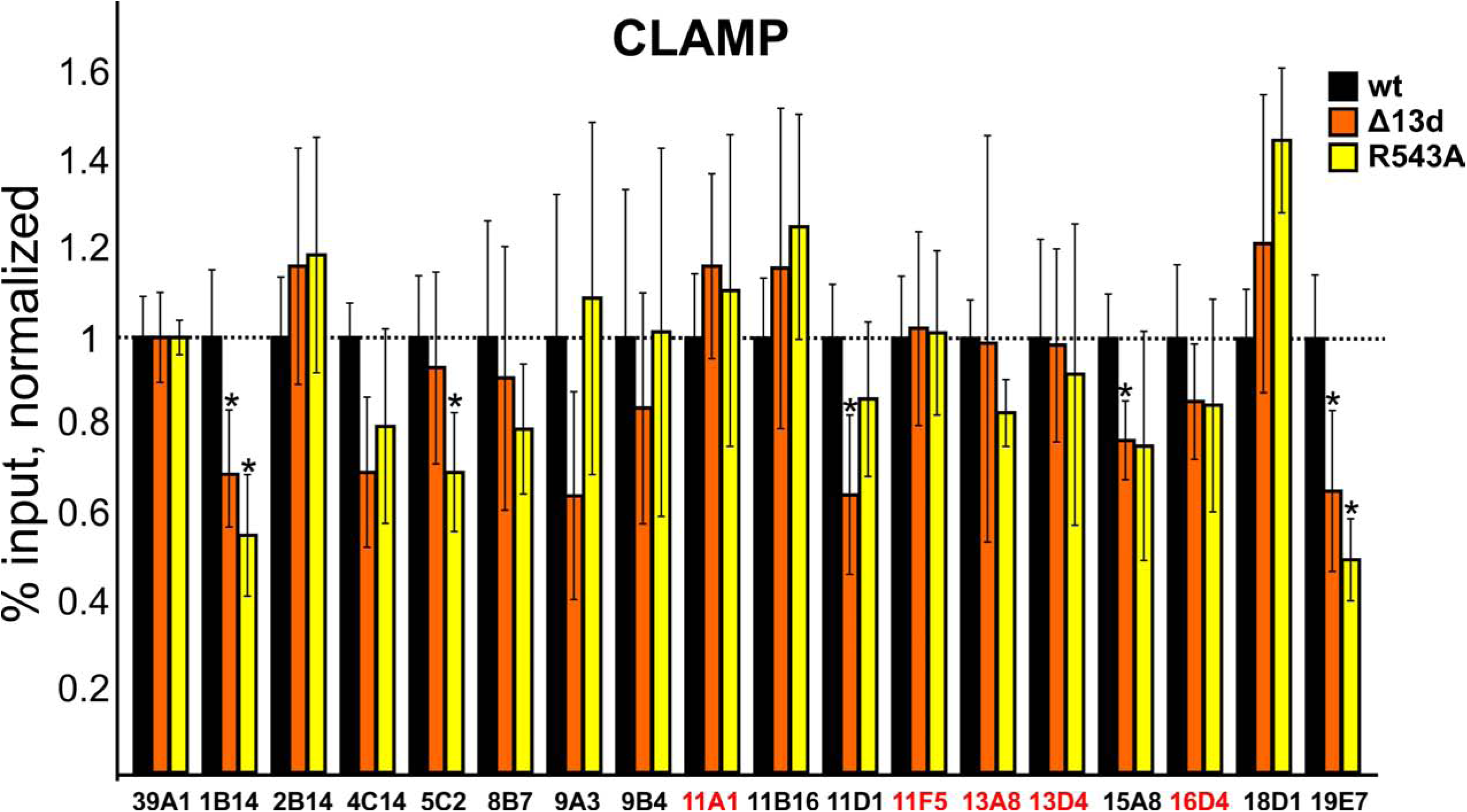
Comparing binding of CLAMP at the different CES and PionX (marked with red) regions in the MSL2-expressing flies at the *msl2*^γ *227*^ background. Histograms show ChIP enrichments at the CES regions on chromatin isolated from male flies expressed different variants (wt, Δ13d, R543A) of MSL2 protein. The results are presented as a percentage of input genomic DNA normalized to corresponding positive autosomal genome regions for CLAMP (39A1), and compared with binding levels in the flies with expressed wild type MSL2 protein (corresponding to the “1” on the scale). Error bars show standard deviations of quadruplicate PCR measurements for three independent experiments. Asterisks indicate significance levels: *p < 0.05.

**Figure EV5.**
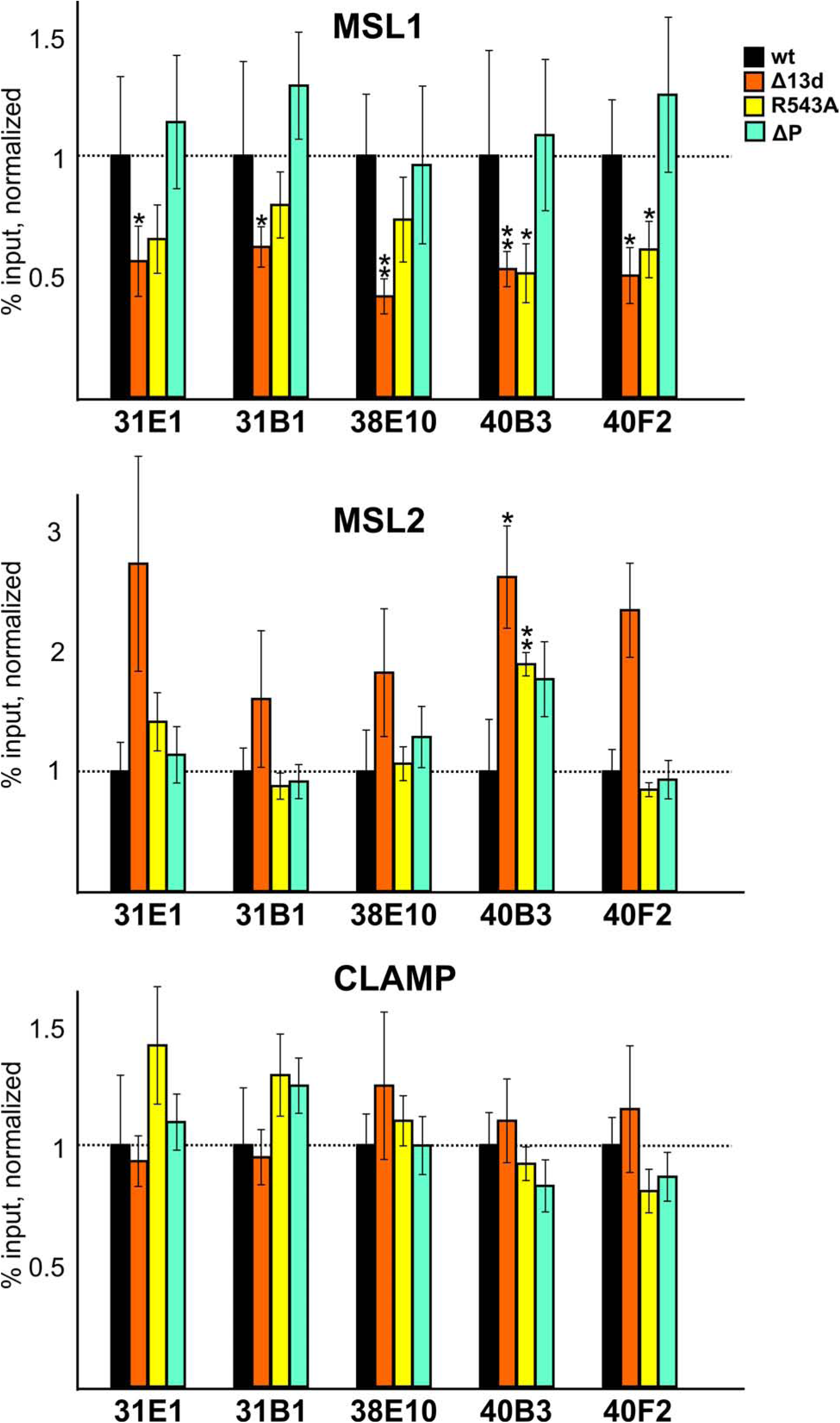
Comparing binding of MSL1 and MSL2 at the different autosomal genome regions in the MSL2-expressing flies at the *msl2*^γ *227*^ background. Histograms show ChIP enrichments at the CES regions on chromatin isolated from male flies expressed different variants (wt, Δ13d, R543A, ΔP) of MSL2 protein. The results are presented as a percentage of input genomic DNA normalized to corresponding positive autosomal genome regions for MSL1(26E3), MSL2 (25A3), and compared with binding levels in the flies with expressed wild type MSL2 protein (corresponding to the “1” on the scale). Error bars show standard deviations of quadruplicate PCR measurements for three independent experiments. Asterisks indicate significance levels: *p < 0.05, **p < 0.01.

## References

Alekseyenko AA, Peng S, Larschan E, Gorchakov AA, Lee OK, Kharchenko P, McGrath SD, Wang CI, Mardis ER, Park PJ, Kuroda MI (2008) A sequence motif within chromatin entry sites directs MSL establishment on the Drosophila X chromosome. Cell 134: 599–609

Bai X, Alekseyenko AA, Kuroda MI (2004) Sequence-specific targeting of MSL complex regulates transcription of the roX RNA genes. The EMBO journal 23: 2853–2861

Bischof J, Maeda RK, Hediger M, Karch F, Basler K (2007) An optimized transgenesis system for Drosophila using germ-line-specific phiC31 integrases. Proceedings of the National Academy of Sciences of the United States of America 104: 3312–3317

Chlamydas S, Holz H, Samata M, Chelmicki T, Georgiev P, Pelechano V, Dundar F, Dasmeh P, Mittler G, Cadete FT, Ramirez F, Conrad T, Wei W, Raja S, Manke T, Luscombe NM, Steinmetz LM, Akhtar A (2016) Functional interplay between MSL1 and CDK7 controls RNA polymerase II Ser5 phosphorylation. Nature structural & molecular biology 23: 580–589

Fauth T, Muller-Planitz F, Konig C, Straub T, Becker PB (2010) The DNA binding CXC domain of MSL2 is required for faithful targeting the Dosage Compensation Complex to the X chromosome. Nucleic acids research 38: 3209–3221

Hallacli E, Lipp M, Georgiev P, Spielman C, Cusack S, Akhtar A, Kadlec J (2012) Msl1-mediated dimerization of the dosage compensation complex is essential for male X-chromosome regulation in Drosophila. Molecular cell 48: 587–600

Ilik IA, Maticzka D, Georgiev P, Gutierrez NM, Backofen R, Akhtar A (2017) A mutually exclusive stem-loop arrangement in roX2 RNA is essential for X-chromosome regulation in Drosophila. Genes & development 31: 1973–1987

Kadlec J, Hallacli E, Lipp M, Holz H, Sanchez-Weatherby J, Cusack S, Akhtar A (2011) Structural basis for MOF and MSL3 recruitment into the dosage compensation complex by MSL1. Nature structural & molecular biology 18: 142–149

Kelley RL, Wang J, Bell L, Kuroda MI (1997) Sex lethal controls dosage compensation in Drosophila by a non-splicing mechanism. Nature 387: 195–199

Kuroda MI, Hilfiker A, Lucchesi JC (2016) Dosage Compensation in Drosophila-a Model for the Coordinate Regulation of Transcription. Genetics 204: 435–450

Kuzu G, Kaye EG, Chery J, Siggers T, Yang L, Dobson JR, Boor S, Bliss J, Liu W, Jogl G, Rohs R, Singh ND, Bulyk ML, Tolstorukov MY, Larschan E (2016) Expansion of GA Dinucleotide Repeats Increases the Density of CLAMP Binding Sites on the X-Chromosome to Promote Drosophila Dosage Compensation. PLoS genetics 12: e1006120

Larschan E, Soruco MM, Lee OK, Peng S, Bishop E, Chery J, Goebel K, Feng J, Park PJ, Kuroda MI (2012) Identification of chromatin-associated regulators of MSL complex targeting in Drosophila dosage compensation. PLoS genetics 8: e1002830

Li F, Parry DA, Scott MJ (2005) The amino-terminal region of Drosophila MSL1 contains basic, glycine-rich, and leucine zipper-like motifs that promote X chromosome binding, self-association, and MSL2 binding, respectively. Molecular and cellular biology 25: 8913–8924

Li F, Schiemann AH, Scott MJ (2008) Incorporation of the noncoding roX RNAs alters the chromatin-binding specificity of the Drosophila MSL1/MSL2 complex. Molecular and cellular biology 28: 1252–1264

Lucchesi JC (2018) Transcriptional modulation of entire chromosomes: dosage compensation. Journal of genetics 97: 357–364

Lyman LM, Copps K, Rastelli L, Kelley RL, Kuroda MI (1997) Drosophila male-specific lethal-2 protein: structure/function analysis and dependence on MSL-1 for chromosome association. Genetics 147: 1743–1753

Maenner S, Muller M, Frohlich J, Langer D, Becker PB (2013) ATP-dependent roX RNA remodeling by the helicase maleless enables specific association of MSL proteins. Molecular cell 51: 174–184

Maksimenko O, Bartkuhn M, Stakhov V, Herold M, Zolotarev N, Jox T, Buxa MK, Kirsch R, Bonchuk A, Fedotova A, Kyrchanova O, Renkawitz R, Georgiev P (2015) Two new insulator proteins, Pita and ZIPIC, target CP190 to chromatin. Genome research 25: 89–99

Meller VH, Wu KH, Roman G, Kuroda MI, Davis RL (1997) roX1 RNA paints the X chromosome of male Drosophila and is regulated by the dosage compensation system. Cell 88: 445–457

Morra R, Yokoyama R, Ling H, Lucchesi JC (2011) Role of the ATPase/helicase maleless (MLE) in the assembly, targeting, spreading and function of the male-specific lethal (MSL) complex of Drosophila. Epigenetics & chromatin 4: 6

Murawska M, Brehm A (2012) Immunostaining of Drosophila polytene chromosomes to investigate recruitment of chromatin-binding proteins. Methods Mol Biol 809: 267–277

Rattner BP, Meller VH (2004) Drosophila male-specific lethal 2 protein controls sex-specific expression of the roX genes. Genetics 166: 1825–1832

Samata M, Akhtar A (2018) Dosage Compensation of the X Chromosome: A Complex Epigenetic Assignment Involving Chromatin Regulators and Long Noncoding RNAs. Annual review of biochemistry 87: 323–350

Soruco MM, Chery J, Bishop EP, Siggers T, Tolstorukov MY, Leydon AR, Sugden AU, Goebel K, Feng J, Xia P, Vedenko A, Bulyk ML, Park PJ, Larschan E (2013) The CLAMP protein links the MSL complex to the X chromosome during Drosophila dosage compensation. Genes & development 27: 1551–1556

Soruco MM, Larschan E (2014) A new player in X identification: the CLAMP protein is a key factor in Drosophila dosage compensation. Chromosome research : an international journal on the molecular, supramolecular and evolutionary aspects of chromosome biology 22: 505–515

Straub T, Grimaud C, Gilfillan GD, Mitterweger A, Becker PB (2008) The chromosomal high-affinity binding sites for the Drosophila dosage compensation complex. PLoS genetics 4: e1000302

Straub T, Zabel A, Gilfillan GD, Feller C, Becker PB (2013) Different chromatin interfaces of the Drosophila dosage compensation complex revealed by high-shear ChIP-seq. Genome research 23: 473–485

Urban J, Kuzu G, Bowman S, Scruggs B, Henriques T, Kingston R, Adelman K, Tolstorukov M, Larschan E (2017) Enhanced chromatin accessibility of the dosage compensated Drosophila male X-chromosome requires the CLAMP zinc finger protein. PloS one 12: e0186855

Villa R, Forne I, Muller M, Imhof A, Straub T, Becker PB (2012) MSL2 combines sensor and effector functions in homeostatic control of the Drosophila dosage compensation machinery. Molecular cell 48: 647–654

Villa R, Schauer T, Smialowski P, Straub T, Becker PB (2016) PionX sites mark the X chromosome for dosage compensation. Nature 537: 244–248

Zheng S, Villa R, Wang J, Feng Y, Becker PB, Ye K (2014) Structural basis of X chromosome DNA recognition by the MSL2 CXC domain during Drosophila dosage compensation. Genes & development 28: 2652–2662

Zhou S, Yang Y, Scott MJ, Pannuti A, Fehr KC, Eisen A, Koonin EV, Fouts DL, Wrightsman R, Manning JE, et al. (1995) Male-specific lethal 2, a dosage compensation gene of Drosophila, undergoes sex-specific regulation and encodes a protein with a RING finger and a metallothionein-like cysteine cluster. The EMBO journal 14: 2884–2895

Zolotarev N, Maksimenko O, Kyrchanova O, Sokolinskaya E, Osadchiy I, Girardot C, Bonchuk A, Ciglar L, Furlong EEM, Georgiev P (2017) Opbp is a new architectural/insulator protein required for ribosomal gene expression. Nucleic acids research 45: 12285–12300

